# Recovery plans need better science to support decisions that allow species to decline in abundance but be recovered

**DOI:** 10.1101/2020.02.27.966101

**Authors:** Ya-Wei Li, Jacob W. Malcom, Judy P. Che-Castaldo, Maile C. Neel

## Abstract

The U.S. Endangered Species Act (ESA) is widely considered the strongest biodiversity conservation law in the world. Part of its strength comes from the mandate to use the best available science to make decisions under the law, including whether to list a species, setting the criteria for when a species can be considered recovered, and determining when those criteria have been met and a species can be delisted. Both biological status and threat factors are considered at each stage of the listing and delisting process. In most cases, conservation science would suggest that species at risk enough to be listed under the Act should be more abundant and secure at delisting than they were at listing. Surprisingly, we identified 130 ESA-listed species that the U.S. Fish and Wildlife Service could consider recovered with fewer populations or individuals than existed at the time of listing. We ask whether their ESA recovery plans present scientific data, rationale, or evidence to support a decline in abundance as part of recovery. We find that almost no plan clearly explains why a decline is allowed. Fewer than half of the plans provide scientific support for a decline in the form of literature references or modeling results. We recommend that the U.S. Fish and Wildlife Service and the National Marine Fisheries Service create a decision support system to inform when species can decline in abundance and still recover, including guidance on (a) the need to explicitly address the declines and (b) the science used to support the decisions.

## Introduction

One of the primary goals of the U.S. Endangered Species Act (ESA) is to recover threatened and endangered species, a purpose that even the law’s critics generally support. There remains robust debate, however, about whether the ESA is effectively achieving that goal. Some members of Congress, for example, regularly condemn the ESA for having taken decades to recover only several dozen species (e.g., Congressional Western Caucus 2019). In response, the federal government has prioritized delisting species that have met their recovery criteria and has highlighted the record number of species recovered in recent years (U.S. Department of the Interior 2019). Often lost in this politically-driven discourse on the effectiveness of ESA recovery is whether the recovery criteria are adequate to ensure a species persists in the long term after it has been delisted under the ESA. Inadequate goals may leave species vulnerable to extinction despite a declaration of recovery. With the growing number of species being recovered, this question becomes increasingly important.

Recovery under the ESA is generally interpreted as the point at which a species is no longer at risk of extinction nor is likely to become so in the foreseeable future (Goble 2009). This interpretation is often recast as a requirement of long-term persistence for a species. Although the time horizon for the future risk of extinction remains undefined in the statute, recent regulatory changes offer some guidance of the term “foreseeable future” (USFWS 2019b). The U.S. Fish and Wildlife Service (FWS) and the National Marine Fisheries Service (NMFS; collectively, the Services) oversee implementation of the ESA, including finalizing recovery plans for listed species. Plans must include estimates of the time and cost of recovery, descriptions of location-specific recovery actions, and criteria for recovering a species. Courts have required the Services to frame those recovery criteria in terms of threats to a species (Fund for Animals v. Babbitt, 903 F. Supp. 96), and the agencies’ recovery planning guidance sets an expectation that recovery plans also include demographic criteria (NMFS 2010). Demographic criteria usually specify population numbers, sizes, trends, and distribution, whereas threat criteria specify the amount that threats must be controlled. The recovery planning guidance explains that demographic criteria should also address the biodiversity principles of representation, resilience, and redundancy (also known as the “3Rs”; Schaffer and Stein 2000, NMFS 2010).

In general, it would be reasonable to expect that the demographic criteria for recovery should specify larger population numbers or sizes than existed at the time of species listing or recovery plan writing. Many species have already declined considerably in abundance by the time they are listed (Neel et al. 2012), and so increases in abundance should lower extinction risk and move those species closer to recovery. However, many recovery plans specified recovery at equal or lower levels of abundance than existed at the time of plan writing or listing (Neel et al. 2012). Some declines were in terms of numbers of populations: 37.1% of 423 species with data required no more populations than existed at listing, and 40.2% of 614 species required no more populations than existed at recovery plan writing. Other declines reflected the number of individual plants or animals: 7.9% of 291 species required no more individuals than existed at listing, and 13.7% of 373 species required no more individuals than existed at recovery plan writing. Even among species that increased in abundance on the path to recovery, most recovery criteria were below levels needed for long-term persistence based on IUCN rankings for numbers of populations and other metrics for viability (Neel et al. 2012). These findings raise serious questions about whether recovery criteria for some species are adequate to ensure long-term persistence. They also raise concern that recovery criteria may be influenced by other factors, such as perceived achievability or sociopolitical risks. Understanding the basis for allowing declines is especially important as an increasing number of species approach their recovery objectives (U.S. Department of the Interior 2019).

There may be scientifically defensible reasons for the Services to specify recovery criteria at lower abundances than existed at the time of plan writing or listing. For example, it may be possible to achieve long-term viability with fewer populations than existed at listing if sacrificed populations are small or otherwise inviable and security of the remaining populations is ensured. Such security can result from alleviating threats such as through habitat protections or habitat management agreements obtained through the ESA (Runge et al. 2007). This tradeoff between threat alleviation and abundance is consistent with the ESA’s focus on threat alleviation in listing and delisting decisions (Friends of Blackwater v. Salazar, 691 F.3d 428). A decline in abundance may even be expected when a species is listed early relative to the full impact of threats. One such example is the listing of the polar bear (*Ursus maritimus*) in the face of climate change well before significant population declines have occurred (USFWS 2008). But no study to date has evaluated whether recovery plans provide justifications for allowing declines in abundance as part of recovery. Without a clear explanation, a recovery plan and the specified criteria may lack scientific credibility and legitimacy with the public. More importantly, such plans drive species closer to extinction and yet still enable the Services to delist them as recovered.

Here, we review the recovery plans that allow species to decline in abundance yet still recover to identify the explanations for the allowed declines. We then evaluate whether those explanations relate to threat amelioration, fulfillment of the 3Rs, or other scientific evidence (e.g., peer-reviewed literature, quantitative or qualitative models). Our analysis focuses not on whether declines would be permitted, but whether the Services recognized the declines and explained them using scientific principles. Finally, we apply the results from this analysis to develop recommendations for improving the specification of recovery criteria and implementation of the ESA.

## Methods

To identify the pool of recovery plans to review, we started with the species that Neel et al. (2012) identified as having recovery criteria that would allow them to be delisted with a decline in abundance (160 species). They evaluated recovery plans through September 2010. We also reviewed all recovery plans finalized between then and September 2015 to identify plans that allowed for declines in abundance. For each species, we recorded the number of individuals and/or the number of populations (a) at the time of listing; (b) at the time of recovery plan writing; and (c) required for delisting. When ranges of abundance were given, we recorded the lower end of the range. This process resulted in identifying four additional species in four plans. For 30 species that Neel and colleagues identified as declining, we found no decline. One reason for this discrepancy is that for species with a range of values for abundance, Neel and colleagues used the upper range of abundances for values at listing and the lower range of abundances at delisting. In this study we used the lowest value of the range for both. In other cases evaluators in this study interpreted the plans differently and reached a different conclusion regarding whether declines were indicated. In still other cases it was determined that there was too much uncertainty to state there would be a decline. And in a few cases, there were typos in the original data that we corrected.

Next, we recorded answers to five key questions that inform understanding of why these species might be allowed to decline in abundance but still be considered to recover:

1. *Does the recovery plan acknowledge the lower abundance in the recovery criteria than levels at the time of listing or recovery plan writing?* For each species, we evaluated this question for abundance in terms of both populations and individuals, and provided the following possible responses: yes, no, N/A. This question mainly addresses whether the plan authors recognized that the recovery criteria stipulates/entails a decline in abundance. N/A indicates that there was no decline for a particular measure of abundance.
2. *If the answer to questions 1 is yes, does the plan specifically state why the lower abundance is adequate for recovery?* This question differs from the first one in that it examines whether the Service explained why it would consider the species recovered even at the lower abundance. For each species, we evaluated this question for both populations and individuals, and provided the following possible responses: explicit explanation, implicit explanation, no explanation, N/A. An “explicit” answer means that the recovery plan provides an explicit explanation, whereas an “implicit” answer means that the plan provides only an indirect explanation but we were able to infer that the authors were addressing the core issue. For example, the 1983 recovery plan for the Indiana bat would consider the species recovered based on protections for 58% of the populations that existed at the time of plan writing. Although the plan does not explicitly state *why* the reduced abundance is adequate to consider the species recovered, it does recognize that the species can be recovered at the reduced abundance if the 58% of populations are protected.
3. *Is threat amelioration one of the reasons given?* Because ESA listing and delisting decisions focus on threat alleviation, we also recorded whether the recovery plan specified that threat amelioration was a reason for the decline. Possible answers were explicit, implicit, or no.
4. *Are the lower abundances based on models, literature review, or other scientific process?* This question addresses the extent to which the lower abundances in the recovery criteria were established based on scientific evidence. The possible answers are quantitative models, semi-quantitative models, verbal models (i.e., logical arguments but without explicit mathematical support), literature references, or no models.
5. *Are abundance levels at recovery supported by an analysis of the levels of resilience, redundancy, and representation?* Existing guidance suggests that demographic recovery criteria should address the 3Rs (NMFS 2010). We recorded whether each of the three R factors were discussed; the possible answers for each factor were explicit, implicit, or no.

We analyzed these data using summary statistics. Specifically we calculated the frequencies and proportions of species that fell into each category of possible answers for each question. We did not perform further analyses because temporal clustering in the date of recovery plans and multispecies plans introduce dependencies that violated assumptions of independence of the data points, and more complex modeling would not add significantly to our understanding of the topic. All analyses were done in R version 3.3 and all of the code and data are included in a package vignette in GitHub (https://github.com/jacob-ogre/decline.recovery) and archived at the Open Science Foundation (https://osf.io/qv425/).

## Results

We identified 130 species that could be considered recovered with a smaller number of populations, individuals, or both compared to the time of listing, recovery plan writing, or both (SI Table 1). Of these, 113 species had at least one type of population decline, and 30 species had at least one type of individual decline (Table 1, Figure 1). Ninety-nine species had decreases in populations but increases in individuals, 17 species had decreases in individuals but increases in populations, and 14 species had decreases in both categories. The large number of species with at least one type of population decline is driven by plants, which comprise 82% of the 130 species (Figure 2).

**Table 1.** Breakdown of 130 species that qualified for our study among four decline categories. Columns and rows sum to over 130 because the categories are not mutually exclusive.

|  | Decline in<br>populations | Decline in<br>individuals | Population and<br>individual declines |
| --- | --- | --- | --- |
| From time of listing | 56 | 8 | 3 |
| From time of plan<br>writing | 109 | 28 | 14 |
| From both listing and<br>plan writing | 52 | 6 | 3 |

**Figure 1.**
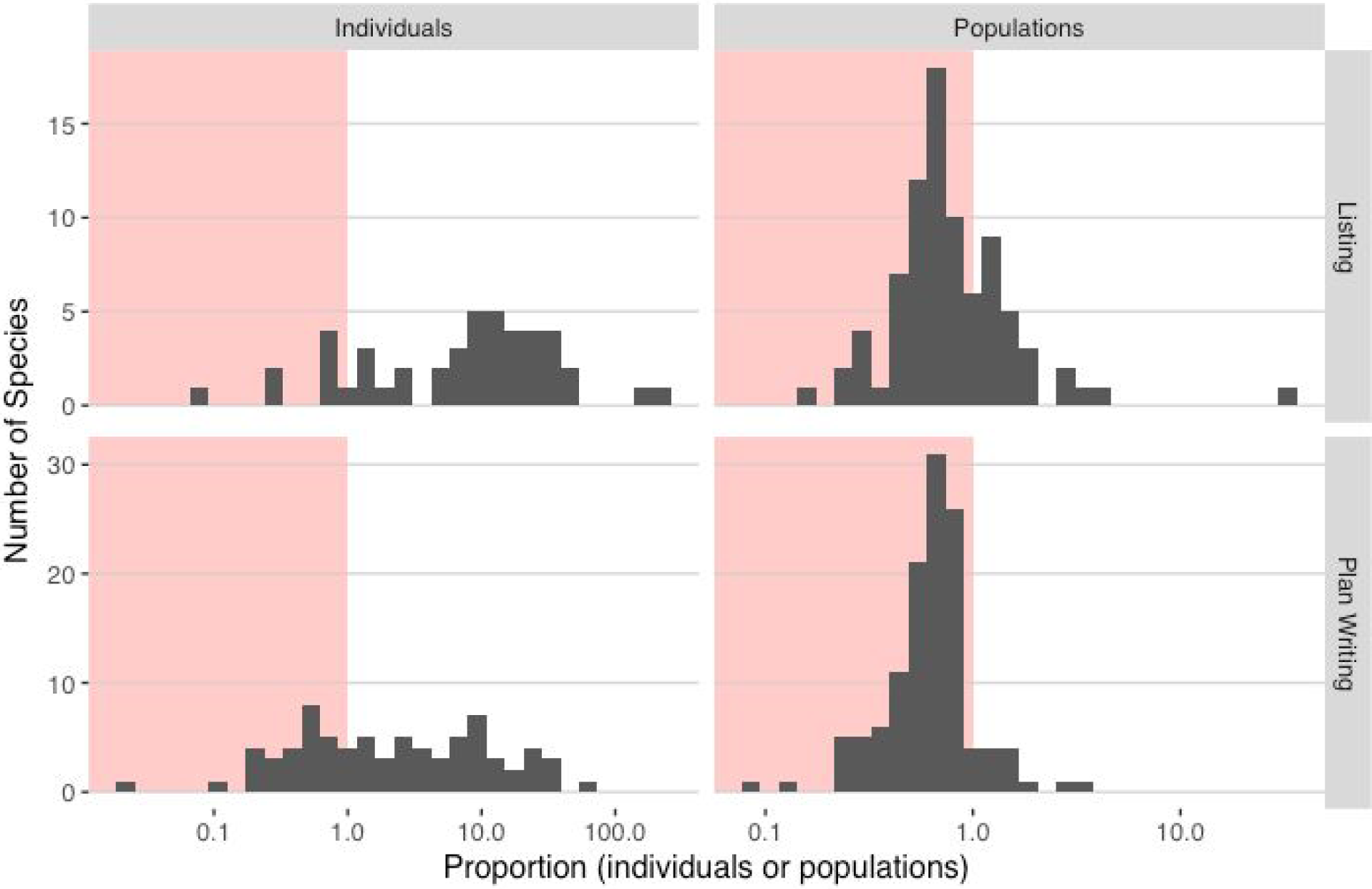
The distributions of the amount of demographic change (proportion) in each of four categories of possible decline. Proportions are calculated as the number of individuals or populations required for recovery divided by the number at listing or at recovery plan writing, so that values < 1 (pink area) indicate allowed declines. Most species qualified based on decline in populations. Each of the 130 qualifying species occurs in at least one panel.

**Figure 2.**
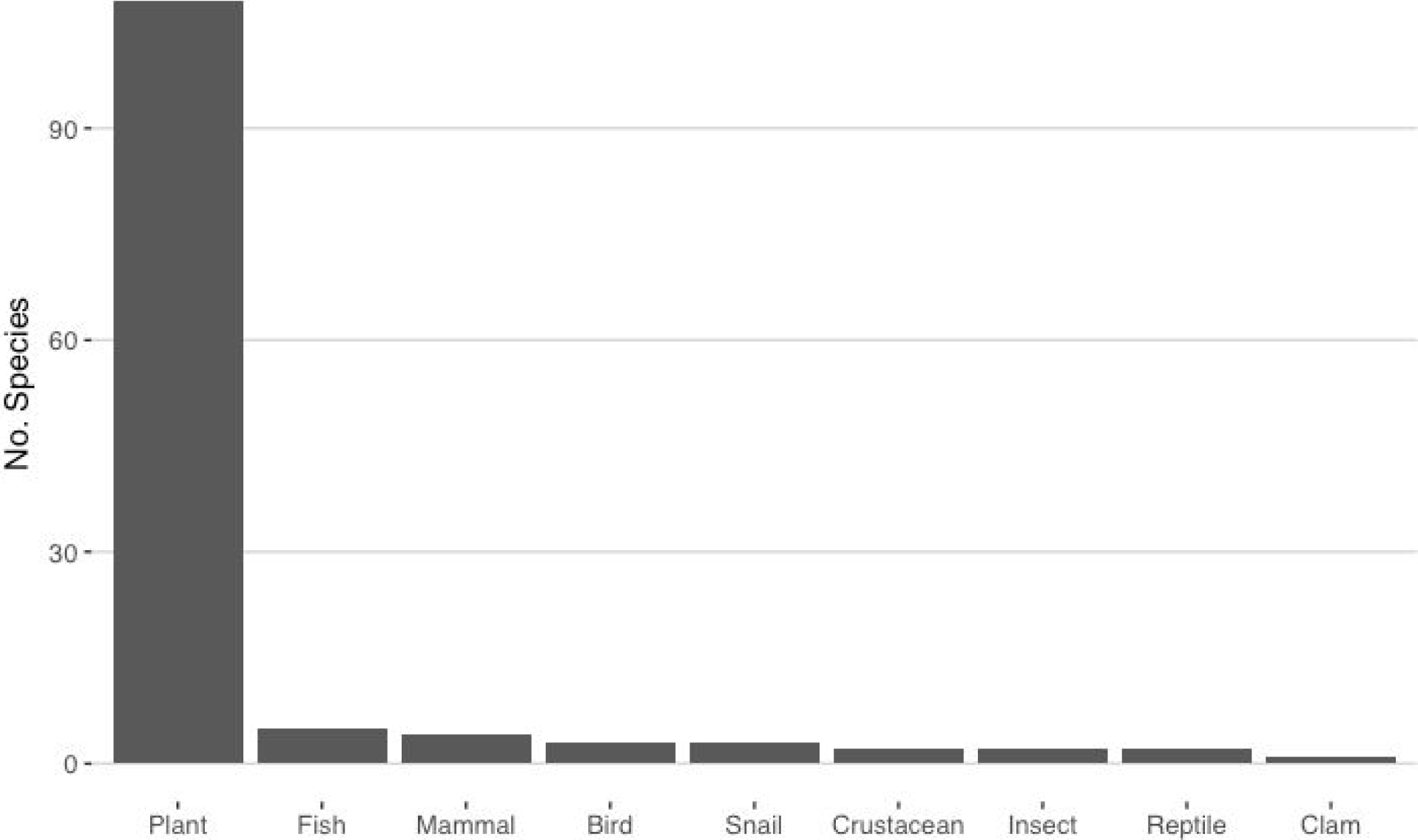
Frequency of taxonomic groups whose species can decline in abundance (populations and/or individuals) and still recover. Plants comprise 82% of the 130 species allowed to decline, but only 57% of Endangered Species Act-listed species.

The plans that allow declines in abundance are not distributed equally across time (Figure 3). Among the 130 species, 96 had plans finalized between 1990 and 2000. This coincides with the increased emphasis on recovery planning in the mid- to late-1990s (Malcom and Li 2018) and extensive use of multispecies plans during that time. These multi-species plans create issues for rigorous statistical analysis of temporal trends. For example, the Maui Plant Cluster Recovery Plan identifies two sets of recovery criteria, one for short-lived species and one for long-lived species. Each of those sets is, ostensibly, adequate to recover plants with those life history characteristics. But because the criteria were not developed individually for each species, the species in that plan are not independent points in our dataset for analysis. Thus, we were unable to identify temporal patterns for allowed declines because of the presence of multispecies plans and the strong temporal clustering in the date of recovery plans in our study.

**Figure 3.**
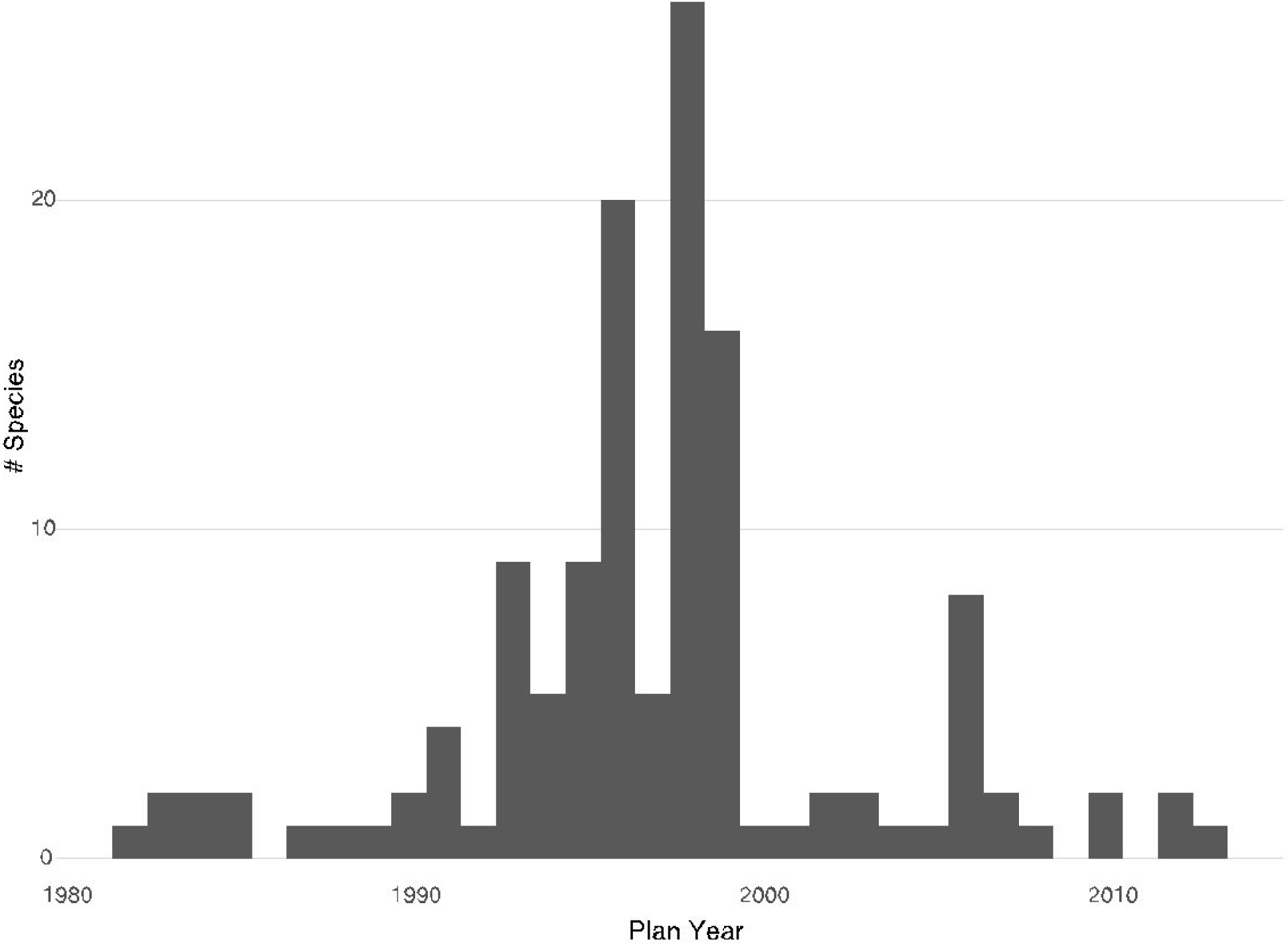
Most recovery plans that allow a species to decline in abundance and still recover were finalized between 1990 and 2000. This period coincides with a significant emphasis on recovery planning by the U.S. Fish and Wildlife Service and the extensive use of multispecies recovery plans.

**Figure 4.**
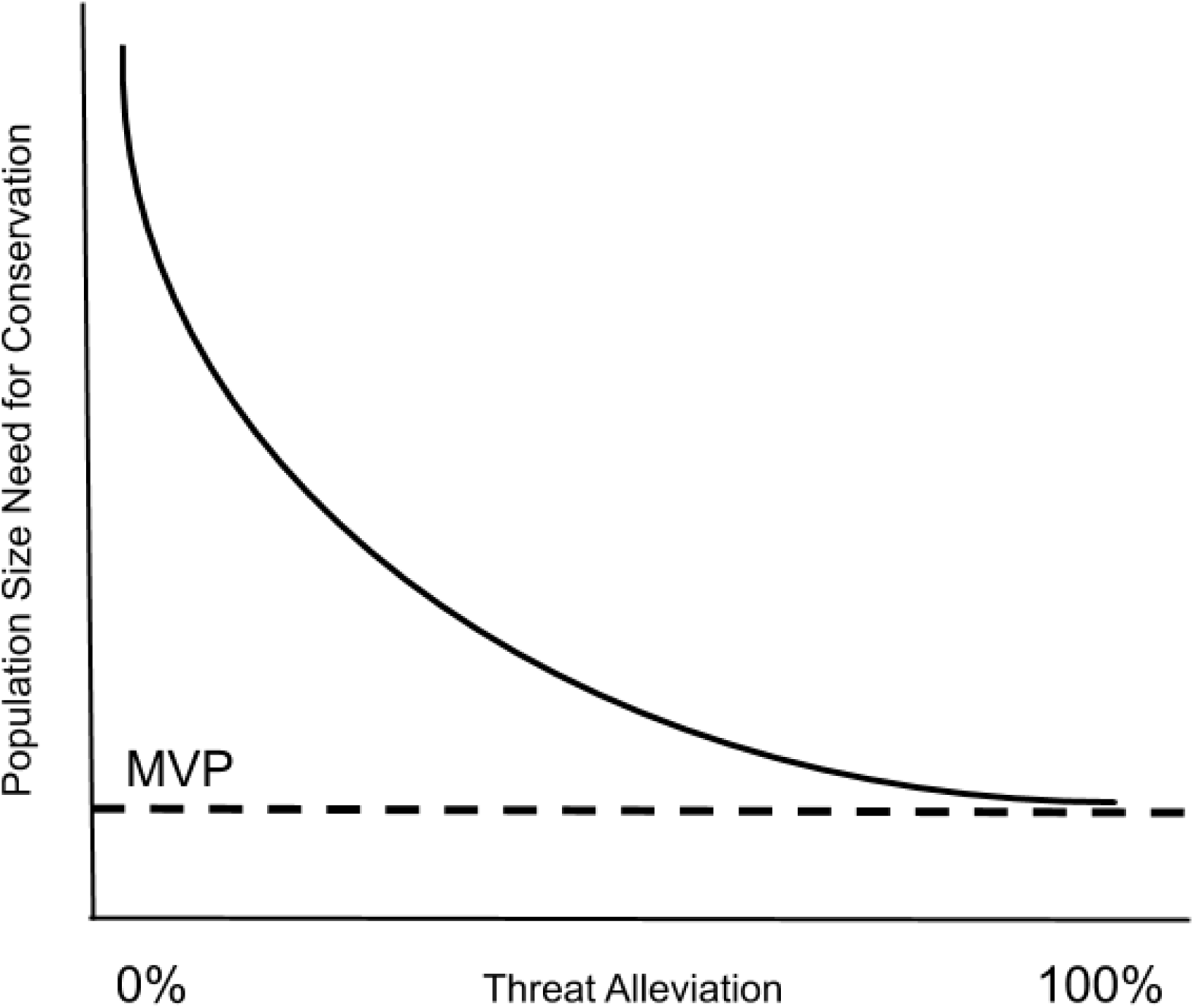
Hypothetical relationships between extent of threat reduction and abundance at recovery needed to maintain long-term viability at a particular level. Both factors are important in ESA delisting decisions because the statute requires the Services to consider biological status and the extent to which each of the five statutorily-identified threat factors have been addressed. MVP = Minimum Viable Population, below which no amount of threat alleviation will allow a species to persist.

### Acknowledgment and explanation for decline in abundance

Most plans do not acknowledge that a species is allowed to decline in abundance and be considered recovered – only 20 acknowledged population declines and 6 acknowledged individual declines (Table 2). For the 20 species for which the population declines were recognized, we then evaluated whether a specific explanation for each decline was given and whether it was explicit or implicit. Only the bog turtle (*Clemmys muhlenbergii*) plan provides an explicit explanation. The plan states that “protection of 185 of the 350 extant bog turtle sites and their populations…has been determined to be appropriate to meet the recovery goal, since protection of this many sites across the species’ range will significantly reduce the species’ risk of extinction due to anthropogenic and non-anthropogenic threats and allow its eventual delisting.” Thus, the plan explicitly addresses why the 185 sites are adequate for recovery. Among the remaining 19 species with a decline in populations, 13 have plans that merely imply an explanation (e.g., Braun’s rocket, see “Threat amelioration” section below) and 6 offer no explanation. For example, the blackside dace (*Phoxinus cumberlandensis*) recovery plan requires the species to inhabit 24 streams before delisting, whereas the species inhabited 35 streams at the time of plan writing. The plan states that “some presently known marginal populations may be lost” but does not explain why 24 populations is adequate for recovery. For species with a decline in terms of numbers of individuals, only five (17%) plans acknowledge the decline, of which four implicitly explain why the recovery criteria are adequate and none does so explicitly.

**Table 2.** Numbers of species with recovery plans that acknowledge a decline in abundance and explain why the lower abundance is adequate for recovery. Some of the columns and rows sum to over 130 because the categories are not mutually exclusive.

|  | Decline in populations | Decline in individuals |
| --- | --- | --- |
| Acknowledgment of decline | 20 | 5 |
| Explicit explanation of adequacy | 1 | 0 |
| Implicit explanation of adequacy | 13 | 4 |
| No explanation of adequacy | 6 | 1 |
| No explanation of decline | 96 | 30 |

### Threat amelioration

Most of the plans we evaluated (*n =* 105; 80%) have no language implying that a decline in abundance is warranted because of threat alleviation through conservation measures described in a recovery plan. The remaining 25 plans (20%) implicitly state threat alleviation as a reason, but offer no explicit explanation for how it was determined the threat alleviation was sufficient. For example, the plan for the Braun’s rockcress (*Arabis perstellata*) sets demographic recovery goals at 20 self-sustaining populations of the plant. At the time of plan writing, there were 28 populations but they were not necessarily secure or self-sustaining. The plan explains that “population levels are declining; eight sites previously known in Kentucky were found to be extirpated in 1996.” To achieve recovery, the 20 populations must be “protected,” which can include “ownership by government or private actions” or “legal dedication or the placement of conservation easements on private land… All protected sites must have management plans or agreements in place that ensure the long-term maintenance of *Arabis perstellata* habitat.” We scored plans like this as providing an “implicit” reason for allowing the decline in abundance. The plan does not explicitly state that a decline is allowed “because” of enhanced conservation measures, but implies a reason for the allowed decline.

### Scientific support for abundance levels at recovery

Plans for 58 (44%) of the species provide no literature reference or verbal, quantitative, or semi-quantitative models to support the population levels required for recovery. For example, the bog turtle plan discussed earlier does not explain how FWS determined that protection of 185 sites would significantly reduce the species’ extinction risk such that it would be deemed recovered. For the remaining 72 species, 46 had plans that provide a literature reference; 58 provide a verbal model; 5 provide a quantitative model; and 2 provide a semi-quantitative model. For example, many plants covered by the Recovery Plan for the Big Island Plant Cluster (e.g., *Silene hawaiiensis*; USFWS 1996) have criteria “… formulated based on recommendations by the Hawaii and Pacific Plants Recovery Coordinating Committee, as well as the International Union for the Conservation of Nature and Natural Resources’ (IUCN’s) draft red list categories (Version 2.2), and the advice and recommendations of various biologists and knowledgeable individuals.” This indicates both literature references and cases in which experts related biological attributes to legal requirements of recovery (i.e., a model of the relationship), but gives no indication of any quantitative or semi-quantitative models used in setting criteria. In contrast, the recovery plan for the Colorado pikeminnow (*Ptychocheilus lucius*) uses quantitative models of multiple processes (e.g., demography and genetics) and scales (e.g., local and metapopulation) to guide criteria development.

### Redundancy, resiliency, and representation

No plan discusses all three Rs explicitly, but most (86) discuss all three Rs implicitly and most (123) discuss at least one of the Rs implicitly or explicitly (Table 3). For example, the copperbelly water snake (*Nerodia erythrogaster neglecta*) recovery plan explicitly requires redundancy of five geographically distinct populations for delisting (e.g., “This ensures redundancy across the landscape to prevent stochastic events from eliminating the entire population of copperbelly”). In contrast, the species’ plan only references the concept of resiliency tangentially and not as part of a guiding framework for setting recovery criteria. Only seven species had plans that did not address representation, resilience, or redundancy at all. These plans were finalized before 1997, at least three years before the 3Rs were formally articulated (Shaffer and Stein 2000).

**Table 3.** Summary statistics of number of species with plans that address resilience, redundancy, and representation.

|  | Explicit | Implicit | None |
| --- | --- | --- | --- |
| Resiliency | 2 | 102 | 26 |
| Redundancy | 4 | 100 | 26 |
| Representation | 12 | 99 | 19 |

## Discussion

In recent years, the public debate on endangered species recovery has often focused on the number and rate of recoveries. But an equally important discussion is whether recovery criteria are scientifically sound (Beissinger 2015, Doak et al. 2015, Neel and Che-Castaldo 2013, Wolf et al. 2015), especially as an increasing number of species approach their recovery goals (U.S. Department of the Interior 2019). The public may view with suspicion government decisions about when to delist a species if there is a decrease in abundance on the path to recovery. A science-based explanation for the decrease should alleviate those concerns and help the Services manage their litigation risk. Our review identified 130 species with lower population and/or individual abundance criteria at the time of recovery than at the time of plan writing and/or listing. Unfortunately, we found that few plans explicitly acknowledge the declines or discuss how they are consistent with recovery, much less provide science-based explanations for the declines.

The near absence of any explicit explanation for the declines highlights a shortcoming in how the Services established recovery criteria for these species. Most recovery plans we reviewed offer no explanation or other information about how the lower abundances would allow for long-term viability such that the species would no longer be at risk of extinction in the near or foreseeable future.

We found that 80% of the 130 plans lack any acknowledgment, much less an explanation, about a tradeoff between abundance and security, even though such tradeoffs may well have been an implied reason for the reduced abundance. We also found that nearly half (44%) of the species examined had plans that offered no scientific literature, verbal, quantitative, or semi-quantitative models to support those abundance levels at recovery. Notably, less than 10% of species had semi-quantitative or quantitative models that informed the recovery criteria, which is comparable to an earlier finding in which ∼13% of recovery plans for plants used or recommended using population viability analyses to set demographic criteria (Zeigler et al. 2011). Our results echo longstanding concerns that the recovery criteria for some species are below levels needed for long-term persistence (Neel et al. 2012) and that some species “would remain in a vulnerable state even if the recovery goals were achieved” (Tear et al. 1995).

We found a strong taxonomic bias to the declines, with plants making up 82% of the species allowed to decline (Figure 2). This bias is consistent with the longer time spent waiting to be listed (Puckett et al. 2016) and substantially less funding (Negrón-Ortiz 2014) for plants than their animal counterparts. Plants are also treated in multispecies recovery plans more often than are animals, and these plans had an outsized effect on the number of species that lacked an explanation for a decline in abundance. Over 50 Hawaiian plant species were covered in five multispecies plant recovery plans (USFWS 1995, 1996a, 1996b, 1997, 1998, 1999), including the 1998 O’ahu Plant Recovery Plan that covers 19 species. None of these five plans offers an explicit explanation for allowing a decline in the recovery criteria. This finding illustrates how shortcomings of a few multispecies plans can propagate to many species. In this case the declines are truly alarming because Hawaiian plants are some of the most endangered species on the list and taxonomic bias in funding precludes careful planning for each species. Such shortcomings can undercut public confidence in the recovery program, a problem that Clark and Harvey (2002) identified in their review of multispecies recovery plans.

In contrast to the paucity of explanations in terms of tradeoffs or scientific models, most plans in this review implicitly or explicitly address at least one component of redundancy, resiliency, and representation, or the 3Rs. In their Interim Endangered Species Recovery Planning Guidance, the Services recommend the 3Rs as one set of principles to analyze in setting recovery criteria (NMFS 2010). This robust discussion of the 3Rs could explain why the Services selected particular recovery criteria without expressly discussing a tradeoff or presenting a biological model. In other words, many Services biologists may have viewed the 3R framework as the primary or sole biological rationale needed to support a particular recovery target. We did not, however, find direct evidence of this in the recovery plans we reviewed, as the discussions of the 3Rs were often not directly linked to the chosen criteria.

We note that the recovery criteria are meant to reflect the *minimum* abundance for delisting, so species may have higher abundances when they are delisted. Nonetheless, our analysis focused on these minimum criteria because the Services could legally delist the species at those levels unless the criteria are revised. Moreover, the criteria are supposed to reflect the best science available at the time of plan writing about the abundance levels needed for long-term viability. Thus, we expected recovery plans to provide science-based explanations about why abundances could decline and how a species could be considered recovered at those lower abundances.

Our analysis identified several opportunities to improve how the Services set recovery criteria:

1. *Develop a decision support system to identify when a species can decline in abundance on the path to recovery and articulate the reasons for the decline*. This recommendation assumes that the Services continue to express demographic recovery criteria in terms of populations, as opposed to the more recent approach where those criteria are occasionally expressed in terms of the probability of long-term persistence (e.g., the polar bear recovery plan; USFWS 2017). In developing this decision support system, the Services should expressly address the potential tradeoff between threat alleviation and abundance: all else being equal, as abundance increases, less threat alleviation may be needed to maintain long-term viability (Figure 2). For the Florida manatee, researchers have quantified how the probability of quasi-extinction varies when key threats to the species (e.g., watercraft mortality) are reduced and when the effective population size used to set the quasi-extinction threshold is changed (Runge et al. 2007). Although the effective population size is not the abundance needed for recovery, the manatee analysis nonetheless shows how population size and threat level can each affect extinction risk. This interplay between abundance and threats likely occurs in every recovery planning effort because the Services consider both species demography and the five statutorily-identified threats in all listing and delisting decisions. The interplay, however, is rarely evident in Services decisions and recovery plans because the agencies have not established a system to objectively determine what level of threat alleviation and population augmentation is needed for recovery. Others have called for the Services to improve the objectivity and consistency of listing and delisting decisions by establishing a quantitative framework to inform those decisions (National Research Council 1995, Doak et al. 2015, Wolf et al. 2015). Our analysis underscores the value of that recommendation because it reiterates the lack of transparency in how the Services arrived at certain recovery criteria. Establishing a framework would allow the agencies to clearly explain when and why certain species might appropriately decline in abundance on the path to recovery. Such a system would also minimize the possibility of Services biologists setting recovery criteria based on the amount of conservation they think is feasible at the time of plan writing rather than the amount needed for long-term species persistence.
2. *Make better use of biological models when establishing demographic criteria*. We recognize that the type of model used will depend on the amount of information available on a species. For data-limited species, literature references and semi-quantitative models may still be feasible approaches. For data-rich species, quantitative models may be needed to satisfy the ESA’s best available science requirement. Given that nearly half of the species in our study had plans with no models and that only seven species had a quantitative or semi-quantitative model, there are ample opportunities in recovery plans to incorporate biological models.
3. *Explicitly discuss and relate the 3Rs to the recovery criteria and actions in every recovery plan*. The 3R framework offers an excellent opportunity for the Services to explain the relationship between the requirements of recovery and core biological principles. This explanation should help encourage broader public support for the recovery criteria the Services set. The framework also serves as a simple checklist for recovery teams and aids in transparency. Through its adoption of the new “Recovery Planning and Implementation” framework (USFWS 2019), which puts the 3Rs at the fore of species status assessments, FWS is likely already on the road to achieving this recommendation. Further, efforts have been made to create measurable and objective criteria for the 3Rs (Wolf et al. 2015, Neel et al. 2014)

Lastly, we note that the 130 species identified in our analysis may not represent all of the species that are allowed to decline on their path to recovery. We were able to identify these species because they have both quantitative demographic recovery criteria, as well as known abundances at the time of listing and/or plan writing. Many listed species do not have measurable recovery criteria or abundance records at different time points, especially plants (Neel et al. 2012, Neel and Che-Castaldo 2013). Recent developments such as the “Recovery Planning and Implementation” framework (FWS 2019) may improve the transparency and consistency of the recovery process, but continued research will be needed to evaluate whether those changes are improving the use of science in informing recovery under the ESA.

## Acknowledgments

We thank Hannah MacDonald and K.C. Stover for assistance in reviewing recovery plans and compiling data. All data and analyses are archived under the Open Science Framework, DOI 10.17605/OSF.IO/QV425.

